## Supplementary Table1 for "A multi-gene predictive model for the radiation sensitivity of nasopharyngeal carcinoma based on machine learning"

| **Variable** | **Overall**(34) | **Radiosensitivity** | | **P value** |
|---|---|---|---|---|
| | | **Rens**(N=23 68%) | **Sens**(N=11 32%) | |
| **Age** | | | | 0.406 |
| Mean(SD) | 46.38(11.7) | 45.26(11.48) | 48.73 (10.98) | |
| Median[Min,Max] | 49.5[26,64] | 49[26,61] | 50[31,64] | |
| **Sex** | | | | 0.942 |
| Female | 9(26.47%) | 6(26.09%) | 3(27.27%) | |
| Male | 25(73.53%) | 17(73.91%) | 8(72.73%) | |
| **Clinical stage** | | | | 0.824 |
| I | 3(8.82%) | 2(8.70%) | 1(9.09%) | |
| II | 4(11.76%) | 3(13.04%) | 1(9.09%) | |
| III | 6(17.65%) | 3(13.04%) | 3(27.27%) | |
| IV | 21(61.76%) | 15(65.22%) | 6(54.55%) | |
| **Pathology** | | | | 0.326 |
| Undifferentiated | 22(64.71%) | 14(60.87%) | 8(72.73%) | |
| Poorly-differentiated | 5(14.71%) | 5(21.74%) | 0(0.00%) | |
| Middle-differentiated | 1(2.94%) | 1(4.35%) | 0(0.00%) | |
| Mix-differentiated | 3(8.82%) | 2(8.70%) | 1(9.09%) | |
| Well-differentiated | 3(8.82%) | 1(4.35%) | 2(18.18%) | |
