## Supplementary Table2 for "A multi-gene predictive model for the radiation sensitivity of nasopharyngeal carcinoma based on machine learning"

| **Characteristic** | **resistant**, N = 1[1] | **sensitive**, N = 3[1] |
|---|---|---|
| Age | | |
| 28 | 0 (0%) | 1 (33%) |
| 34 | 0 (0%) | 1 (33%) |
| 42 | 1 (100%) | 0 (0%) |
| 71 | 0 (0%) | 1 (33%) |
| Sex | | |
| female | 0 (0%) | 2 (67%) |
| male | 1 (100%) | 1 (33%) |
| Clinical Stage | | |
| II | 1 (100%) | 1 (33%) |
| III | 0 (0%) | 2 (67%) |
| Pathology | | |
| Undifferentiated non-keratinized | 1 (100%) | 3 (100%) |

[1]n (%)
