## Supplementary figures and images for "A multi-gene predictive model for the radiation sensitivity of nasopharyngeal carcinoma based on machine learning"

### Supplementary Figure1

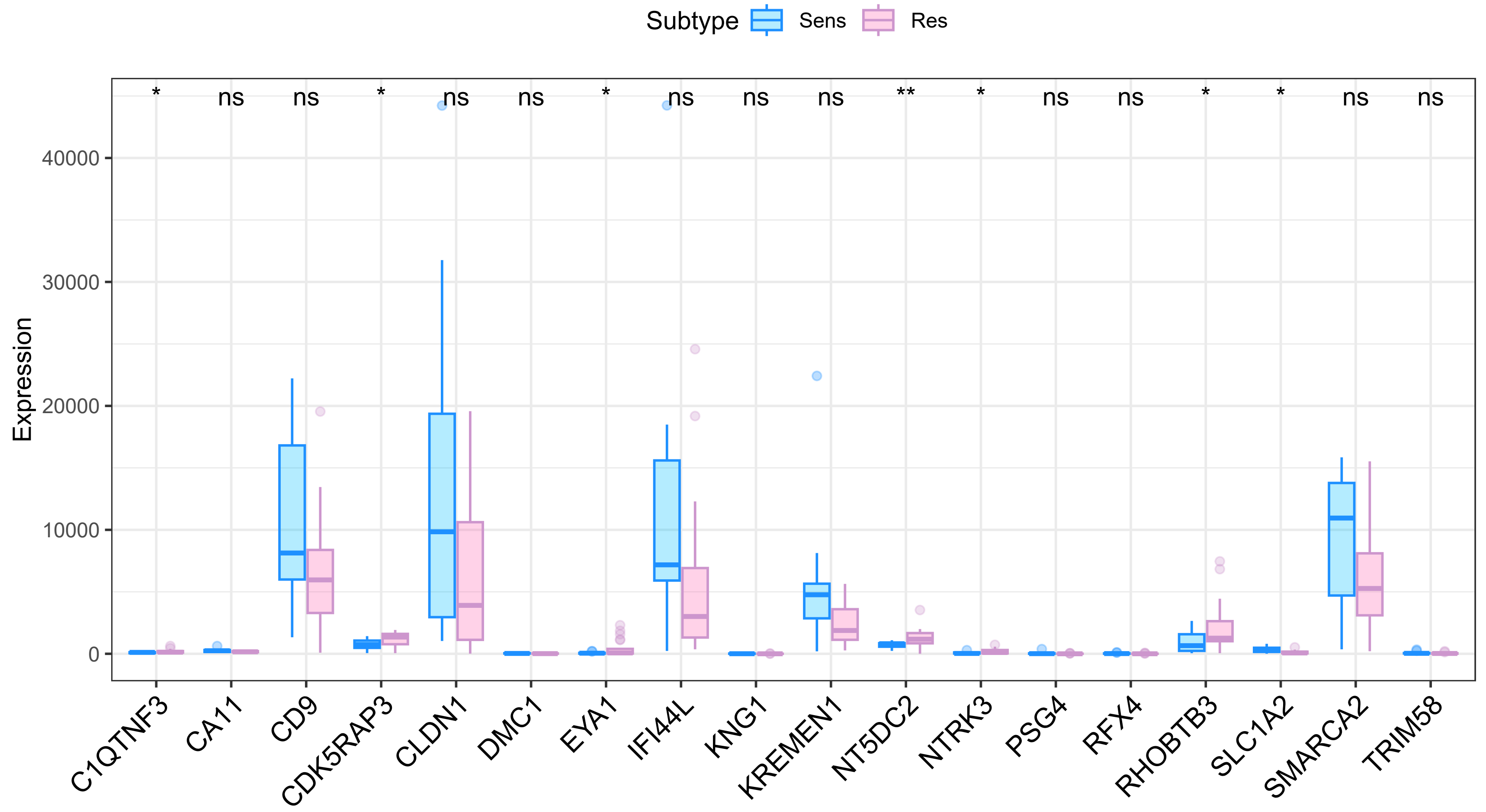

### Supplementary Figure2

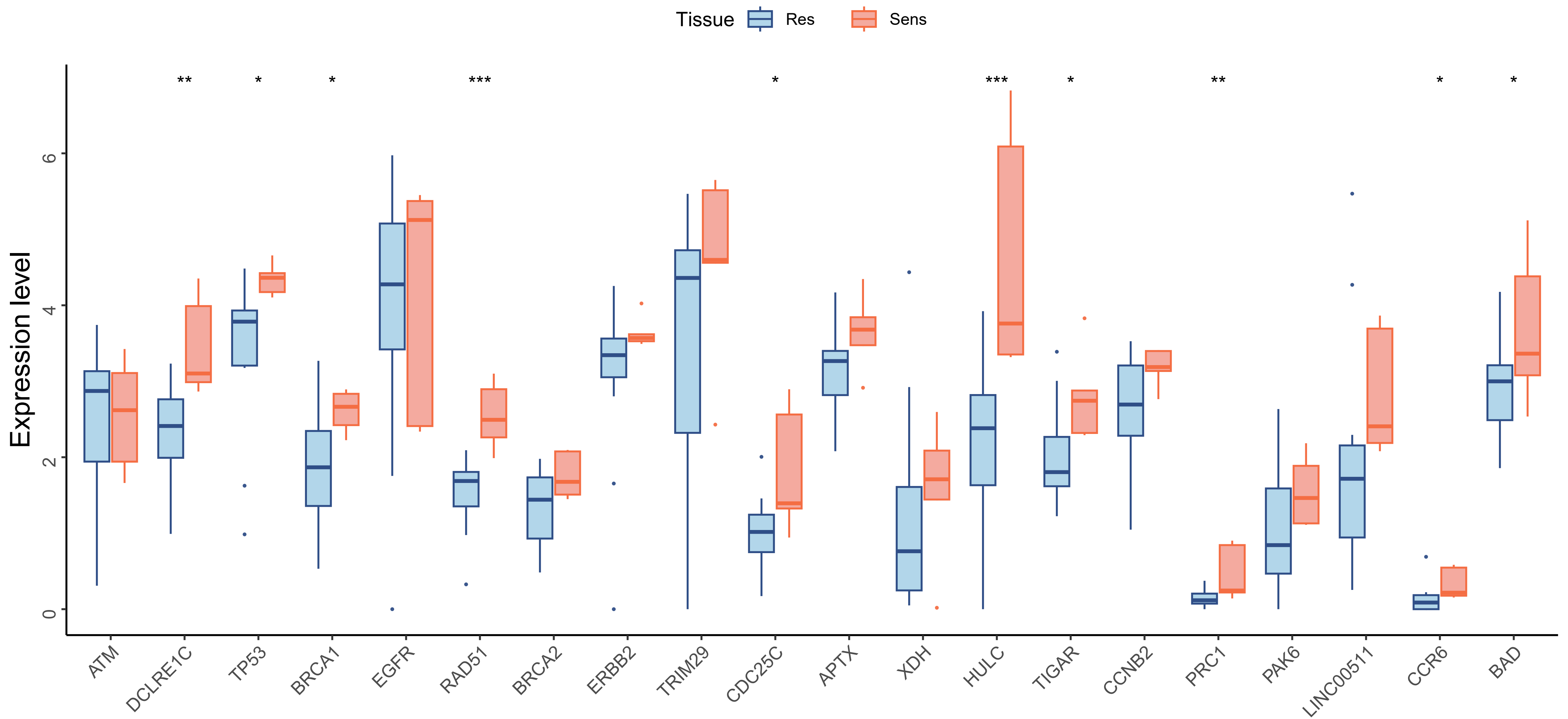
